# Leveraging Multi-Modal Feature Learning for Predictions of Antibody Viscosity

**DOI:** 10.1101/2024.08.30.610550

**Authors:** Krishna D. B. Anapindi, Kai Liu, Willie Wang, Yao Yu, Yan He, Edward J. Hsieh, Ying Huang, Daniela Tomazela

**Affiliations:** Protein Therapeutics, Gilead Sciences; Research Data Sciences, Gilead Sciences

**Keywords:** Multimodal Feature Learning, Antibody Viscosity Prediction, Machine Learning, Bioinformatics

## Abstract

The shift towards the subcutaneous administration route for biologics therapeutics has gained momentum due to its patient-friendly nature, convenience, reduced healthcare burden, and improved compliance compared to traditional intravenous infusions. However, one of the potentially significant challenges with this transition is managing the viscosity of the administered solutions. High viscosity can pose substantial development and manufacturability challenges, directly impacting the patient by increasing injection time and pain at the injection site. Moreover, high viscosity formulations can prolong residence time at the injection site, affecting absorption kinetics and potentially altering the intended pharmacological profile and therapeutic efficacy of the biologic candidate. This publication explores the application of a multimodal feature learning workflow for predicting the viscosity of antibodies in therapeutics discovery, integrating multiple data sources such as sequence, structural, physicochemical properties, and embeddings from a language model. This approach enables the model to learn from various underlying rules, including physicochemical rules from molecular simulations and molecular protein evolutionary rules by large, pre-trained deep learning foundation models. By comparing the effectiveness of this approach against other selected published viscosity prediction methods, this study offers insights on their intrinsic viscosity predictive potential and usability in therapeutics antibody early development pipelines.

## 1 Introduction

The viscosity of a biologic candidate plays a critical role in the feasibility of the therapeutic drug, particularly in the context of subcutaneous administration, where higher concentration preparations are required given the low injection volume required (1-2 mL). A biologic candidate with high viscosity can face development and manufacturability challenges and directly impact patients with increased injection time and pain in the injection site. In addition, high viscosity formulations can prolong the drug residence time at the injection site affecting absorption kinetics and potentially altering the intended pharmacological profile and therapeutic efficacy of the biologic candidate. In early stages of biologics discovery, viscosity read-outs are, in general, not feasible, given the large amount of protein reagent required. Therefore, tools that enable early assessment and consideration of viscosity during sequence selection are imperative to ensure a streamlined drug discovery process, and future enhanced patient experience and clinical outcomes.

Despite significant advancements in machine learning (ML) across various scientific fields, its application in predicting mAb viscosity has been notably limited^1–5^. This limitation primarily stems from the scarcity of high-throughput experimental data in the early phases of discovery and the complexity of capturing the intricate molecular interactions that influence viscosity at elevated concentrations. Additionally, many existing predictive models struggle with issues related to accessibility, computational complexity, and a lack of interpretability, which significantly diminishes their practical utility.

The existing models for predicting monoclonal antibody (mAb) viscosity consider a limited set of features, which are known to impact viscosity either directly or indirectly. For instance, the Sharma model estimates viscosity by considering the Fv net charge, charge asymmetry, and hydrophobicity index based on the protein sequence^5^. Another model calculates a viscosity score using surface charge from molecular dynamics (MD) simulations, referred to as the Surface Charge Model (SCM)^1^. Additionally, a recent study from Pfizer introduced a CNN-based deep learning model (PfAbNet), which is trained on 3-D electrostatic surface potential (3D-ESP) data^4^. While these models aim to estimate the same property—viscosity—their limited feature sets may lead to discrepancies in predictions. In this work, we address the issue of insufficient features by expanding the feature set to include both structural properties derived from modeling via Molecular Operating Environment (MOE) and embeddings from a language model.

Building on this foundation, we present a novel multimodal machine learning framework (MMF) designed to predict the viscosity of mAbs more accurately. Our approach combines protein structure and physicochemical properties derived from molecular models (MOE) with embeddings from two large language models trained on millions of protein sequences: a general protein model (ESM-2)^6^ and an antibody-specific model (AbLang)^7^. These models are fine-tuned to capture the structural and biophysical properties of proteins, transforming them into numerical vectors known as embeddings.

The use of such embeddings as features for training models related to antibody developability has recently been explored by a few research groups^8,9^. Our model leverages these protein-focused embeddings alongside molecular descriptors from the Molecular Operating Environment (MOE). We trained the model using publicly available viscosity datasets, fine-tuning it to achieve superior generalizability across diverse sequences compared to existing models. Furthermore, we demonstrated that our model outperforms others at a practical viscosity cutoff value of 20 cP. In contrast, other models often require adjusting the cutoff value based on the ROC curve, which can be significantly higher than 20 cP when the dataset is biased towards high viscosity samples.

Our MMF approach has identified novel features correlated with viscosity, such as helicity, radius of gyration, and the isoelectric point (pI) of the sequence, in addition to previously known factors like patch hydrophobicity and charge asymmetry. This approach enhances our understanding of model behaviors and establishes a foundation for improving model reliability and interpretability. Consequently, our work advances the strategic use of machine learning, specifically protein language model embeddings, in the development of monoclonal antibody (mAb) therapeutic candidates.

## 2 Methods

### 2.1 Public Datasets

The initial model development was performed on the publicly available sequences and viscosity information from two sources 1) Ab21: List of 21 IgG1 antibodies that are approved by FDA for therapeutic use. 2) PDGF38: List of 38 anti-PDGF sequences by Rai et al. The heavy and light chain sequences of all the antibodies, along with their experimental viscosity at a concentration of 150 mg/mL, were obtained from respective previous works^4,10^.

### 2.2 Internal Sequences

For the internal sequences, we identified 15 IgG1 sequences, all buffer-exchanged to 20 mM Histidine (pH 6.0) and concentrated to 150 mg/mL. All sequences used are wild-type IgG1. They are all reconstructed into a human IgG1 backbone with kappa (15) or lambda (1) light chain. The germline of the sixteen mAbs are very diverse as showed in **SI Table. 1**, with a total of 13 different V_H_ germlines and 11 different V_L_ germlines.

### 2.3 Production of Molecules

All internal WT IgG1 mAbs were expressed in either Exp293, ExpiCHO, or CHOZN cell lines. Protein A chromatography was used as capturing step, followed by polishing with size exclusion chromatography (SEC) or cation exchange (CIEX) chromatography as needed to achieve >95% purity as measured by analytical SEC (aSEC). Samples were dialyzed against their preferred buffer, filtered through 0.2um filter, aliquoted, and stored at -80°C.

### 2.4 Viscosity Measurement

All samples were buffer exchanged into 20 mM Histidine, pH 5.8 and concentrated to >150 mg/ml using a 10 kDa MWCO filter plate on Big Tuna (Unchained Labs, TUNA-0104). Concentration was measured in triplicate, at 280 nm using a Stunner instrument (Unchained Labs, 901006) and purity/aggregation confirmed to be >95% by aSEC. Samples were adjusted to 150 mg/ml before viscosity measurement in triplicates using a VROC Initium one plus system (Rheosense, INI-H-1100) at room temperature.

### 2.5 Antibody Structural Modeling

MOE 2022.02 was used to build homology models from Fv sequences (N=1). A custom script was used for structure generation/preparation and a series (N=90) of protein properties were calculated at pH 5.8.

## 3 Model Development

The Multi-Modal Feature (MMF) model incorporates features from three primary categories:

1. Sequencee-derived descriptors: Previously validated for viscosity prediction and include parameters (q, qsym, and HI) from the Sharma model.
2. Protein structure descriptors: Calculated using MOE (Molecular Operating Environment) software, based on Fv sequence-derived structure models generated through native homology modeling. Examples include surface patch areas (hydrophobic, positive, negative), dipole moment, and helicity. In total, 90 descriptors were computed at the protein level using MOE’s internal pipeline.
3. Protein embeddings: Extracted from pre-trained large protein language models, specifically AbLang and ESM-2 (Evolutionary Scale Modeling 2).

For protein embeddings, we leveraged the strengths of both the language models and their embedding representations. AbLang^7^ is a transformer-based antibody-specific language model with approximately 85 million parameters, trained on the Observed Antibody Space (OAS) database, which comprises 14 million heavy chains (HC) and 0.19 million light chains (LC). We extracted 768-dimensional vector embeddings for both HC and LC variable domains from AbLang. ESM-2^6^, on the other hand, is a general in-purpose, pan-protein pre-trained foundation model based on the BERT transformer architecture^11^, trained on UniRef50. We utilized the 650 million parameter version of ESM-2 to extract 1280-dimensional embedding vectors, following the default guidelines provided in the GitHub repository. For ESM-2, HC and LC variable domain sequences were concatenated using a G4S linker.

While other popular structure modeling software, such as Schrodinger, could potentially provide additional structural descriptor features, we focused solely on MOE in this study. This decision was made to avoid potential redundancy, as features from both MOE and Schrodinger often cover similar aspects of protein models.

The MMF models were trained as Random Forest Regression model to predict the measurement of viscosity assay. The training and evaluation of the models was conducted with a total of 59 mAb molecules from public space plus a cohort of internal 20 mAbs molecules. For ease of comparison with previous models’ performance, 59 mAbs were split into two groups based on the origin of the two datasets, (i) 38 anti-PDGF antibody variants (PDGF 38 set) that lacks sequence and germline diversity but a wide range of viscosity measurement at 150 mg/mL; (2) 21 FDA-approved antibodies (Ab21) with higher sequence and germline diversity compared to PDGF 38 set. Exploratory analysis via t-SNE analysis with all 2909 features from the 59 mAbs suggests that combining MOE descriptors with language model embeddings can further differentiate molecules therefore has the potential to enhance model performance if utilized in training (**SI Fig. 1 A,B**).

**Figure 1.**
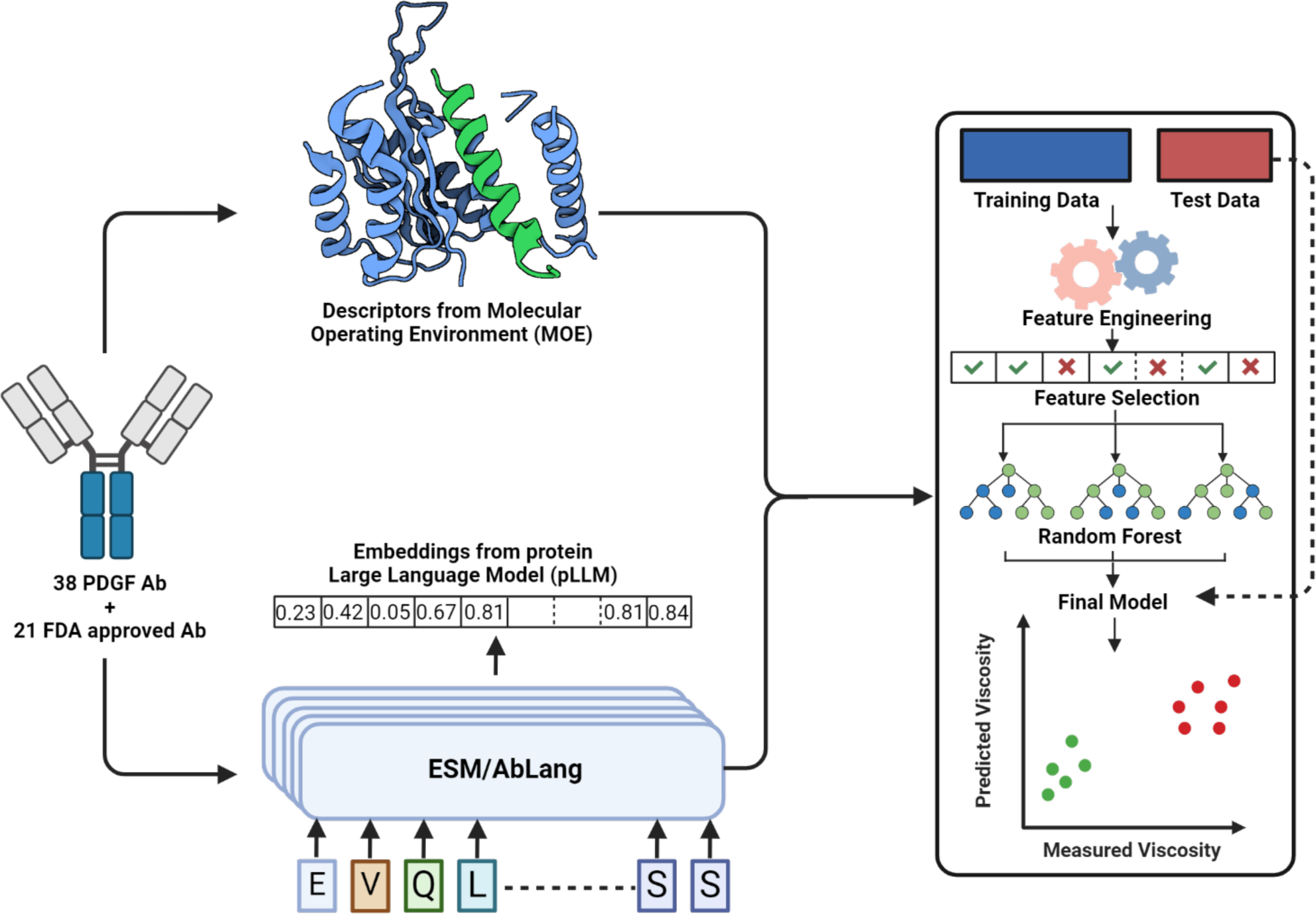
Workflow for the implementation of Multimodal Feature Learning (MMF). Descriptors from MOE are combined with embeddings from protein language models (pLMs). The data is then split into train and test sets, followed by feature engineering (to generate additional descriptors), feature selection and model training. The final model is evaluated on the held-out test set.

Three separate MMF models were trained from scratch namely (1) MMF-PDGF38 (trained on PDGF 38 set), (2) MMF-Ab21 (trained on Ab 21 set) and (3) MMF-LOOCV (trained on PDGF38 and Ab 21 set using leave-one-out cross-validation data split where 58 mAbs including 38 mAbs from PDGF38 set plus 20 mAbs from Ab 21 set were used for training while each of Ab 21 set was held-out as the test set).

Before training, to avoid potential overfitting due to large number (2909) of features compared to sample size, feature selection steps were conducted by (i) spearman rank order correlation ranking; (ii) AUC score ranking - further features were selected with using a base model (random forest regressor with mean absolute error criterion) consisting of hand selected features known to predict viscosity (e.g. sequence descriptors from the Sharma model and three protein descriptors from Makowski et al.^12^ classifier model); (iii) recursive feature selection; and (iv) exhaustive feature selection. In addition, engineered features such as baseline classifier model predicted class label were included in final feature list before steps iii and iv. Next, all features including ranking selected features, base model selected features and engineered features were trained on RF regressors. Then the performance of each model was evaluated on the corresponding test set.

## 4 Results

Two approaches were adopted to compare MMF models with previous models. First, we frame the prediction problem as regression and compare the performance in correlation to determine how close the models can reproduce the specific ground truth value in the data. Second, to further evaluate the model’s applicability in selecting molecules in development pipeline, we frame the problem as classification by classifying each antibody as high or low viscosity based on the predicted value and a practical arbitrary threshold adopted internally (20 cP at 150 mg/mL). As illustrated in Table 1, the MMF model surpasses the current state-of-the-art PfAbNet model when trained on PDGF38 and tested on Ab21 (tab0.78 ± 0.096 vs. 0.75 ± 0.129) as well as in the PDGF38+Ab21 leave-one-out cross-validation (LOOCV) (0.77 ± 0.068 vs. 0.71 ± 0.145) (Pairwise t-test p-values:1.84e-76). Additionally, the MMF model demonstrates nearly equivalent performance to PfAbNet when trained on Ab21 and tested on PDGF38 (0.79 ± 0.051 vs. 0.80 ± 0.053). The p-value comparison for pairwise correlation is shown in **SI Fig.2** and **SI Table 2**. For the evaluation based on classification, we compared the accuracy using both a practical viscosity cutoff point (20 cP) and an optimal operating point (OOP). While PfAbNet demonstrates a classification accuracy of 0.76 on the Ab21 dataset, this performance requires adjusting the cutoff value for differentiating high and low viscosity samples from 20 cP to 72 cP, in accordance with the OOP of the AUROC curve (**Fig. 3A,B**). When using the practical cutoff of 20 cP, PfAbNet’s classification accuracy decreases to 0.67 (**SI Table 2A**). In contrast, the MMF model achieves superior performance, maintaining a consistent cutoff of 20 cP for both the practical threshold and the OOP based on AUROC (**Fig. 3C,D**). Additionally, the MMF model also achieves a superior classification accuracy of 0.9 at the 20cP cutoff (**SI Table 2B**).

**Figure 3:**
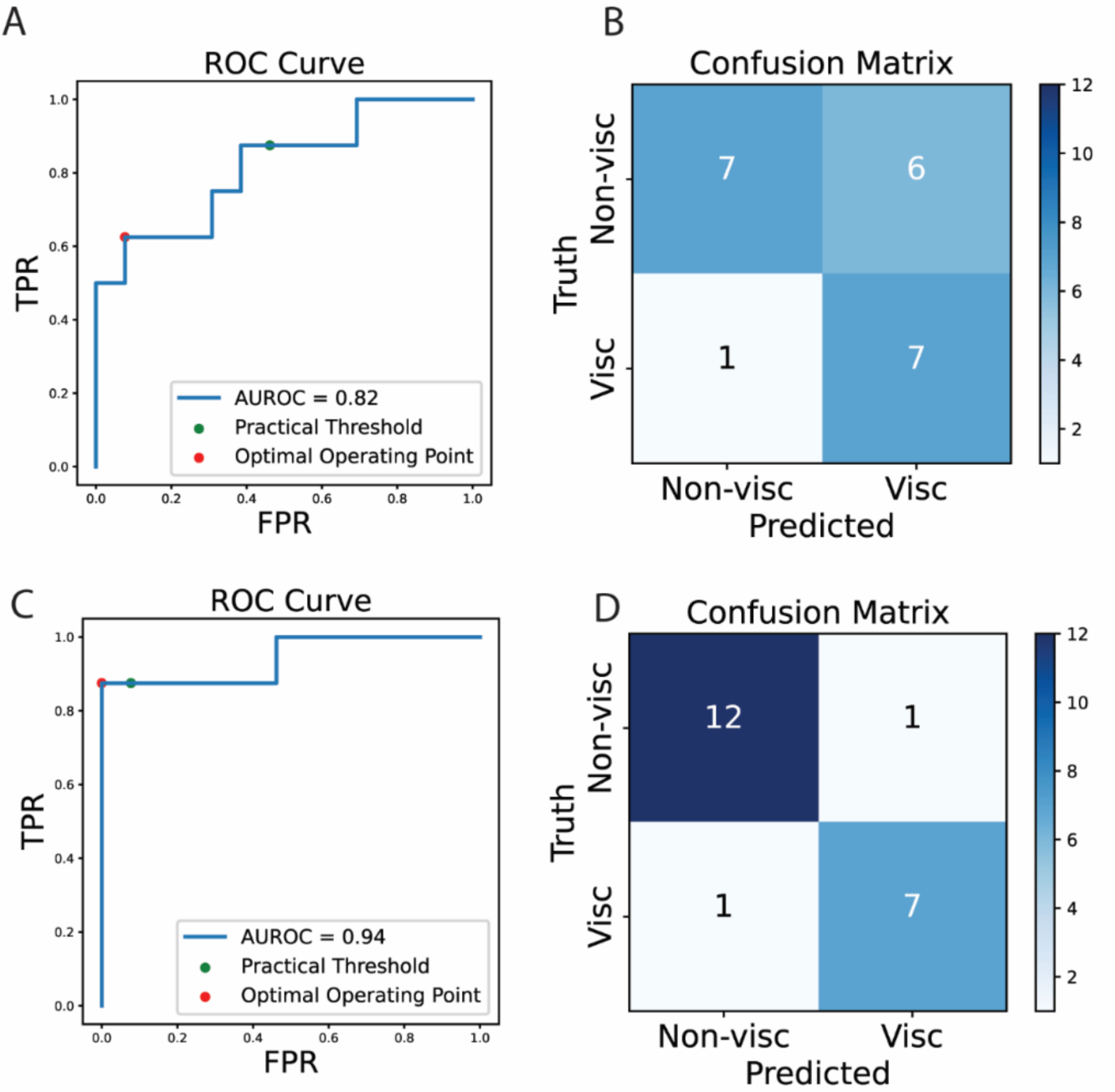
A) ROC curve for PfAbNet with optimal operating point (OOP) significantly different from practical threshold, B) Confusion matrix for PfAbNet at practical threshold, C) ROC curve for MMF with OOP similar to practical threshold and D) confusion matrix for MMF at practical threshold.

**Table 1:** Comparative analysis of multimodal feature learning models against previous existing methods. The Spearman rank order correlation corresponds to the model performance in regression task while the ROC-AUC represents the classification model performance.

|  |  | Spearman rank order correlation |  |  |  |
| --- | --- | --- | --- | --- | --- |
| Training | Test | Sharma | SCM | PfAbNet | MMF |
| Ab21 | PDGF38 | 0.67 | 0.75 | 0.80 | 0.79 |
| PDGF38 | Ab21 | 0.65 | 0.49 | 0.75 | 0.78 |
| PDGF38+Ab21 (LOOCV) | Ab21 | 0.55 | 0.49 | 0.71 | 0.77 |
|  |  | ROC-AUC |  |  |  |
| Training | Test | Sharma | SCM | PfAbNet | MMF |
| Ab21 | PDGF38 | 0.73 | 0.8 | 0.87 | 0.88 |
| PDGF38 | Ab21 | 0.78 | 0.63 | 0.82 | 0.91 |
| PDGF38+Ab21 (LOOCV) | Ab21 | 0.74 | 0.63 | 0.82 | 0.88 |

For further internal validations, we use an internal set of antibodies that spread over different germlines and sequence space as true test set. MMF models performed at similar or better level in spearman rank order correlation and displayed less skewedness due to unbalanced training data than earlier models in predicting high viscosity molecules. To our knowledge, this represents the first attempt applying predictive models for viscosity trained with the combination of features from sequence descriptors, structure descriptors and foundation model embeddings.

## 5 Discussion

### 5.1 Improvement in Interpretability and usability of model predictions

Several in-silico models have been developed to predict viscosity, each with its own limitations regarding the features they consider. Most models focus on a limited set of features or those already known to impact viscosity, such as hydrophobicity-based or charge-based features. For instance, the Sharma model, one of the earliest approaches, utilizes three features: net charge, Fv charge symmetry (FvCSP), and hydrophobicity index (HI). However, it is constrained by its dataset, which includes only 14 different monoclonal antibodies (mAbs) and relies solely on sequence-based parameters. This simplistic approach fails to capture the complex interactions that govern viscosity, including structural conformation and higher-order interactions. Another widely used surrogate model, the spatial charge map (SCM), predicts high-viscosity antibodies using electrostatic properties derived from antibody sequences. Despite its utility, the SCM score does not account for other properties affecting viscosity, such as hydrophobic patch area and solvent-accessible surface area. Additionally, the SCM approach lacks a consistent cutoff score for categorizing molecules and serves as a surrogate score rather than providing actual viscosity predictions, relying heavily on accurate homology predictions. The more recent PfAbNet model employs a 3D convolutional neural network (3D-CNN) that uses the electrostatic potential surface grid (ESP-grid) of the antibody variable region as its primary input. While charge-related properties like ESP and SCM correlate with antibody viscosity, relying on a single 3D feature can be limiting for accurately estimating viscosity.

To explore previously unconsidered features and capture novel relationships between protein sequences and biophysical properties, we implemented an innovative approach that combines over 90 calculated properties from the Molecular Operating Environment (MOE) with embeddings from two protein large language models (pLLMs), ESM-2 and AbLang. Protein large language models can mathematically capture the complex dependencies between amino acid residues and biophysical properties by learning intricate patterns and co-evolutionary constraints within the vast sequence space of natural proteins. Our approach, termed multimodal feature learning (MMF), aims to develop a predictive model that not only identifies features from the known repertoire provided by MOE but also leverages the complex, hard-to-discern patterns that influence protein viscosity. This unique approach addresses the black-box nature of large language models, resulting in a more interpretable model.

### 5.2 Identification of informative features by exploratory analysis combining domain knowledge

To rationalize the Multi-Modal Fusion (MMF) model, we conducted a pairwise analysis of MOE features and pLLM embeddings to understand the practical relevance of the selected embeddings. By ranking MOE features based on their frequency in the top 20 correlations with embeddings, we identified 8 groups of descriptors, comprising a total of 38 MOE features (**Fig.4, Appendix I**). Notably, several embedding-correlated features, such as patch hydrophobicity (patch_hyd), Fv charge asymmetry (Fv_chml), and patch positive charge (patch_pos), align with top features identified in other viscosity prediction models^13–16^. Given the complex, non-linear and black-box nature of pLLM embeddings and how they contribute towards the final model performance, we only tried to infer from the absolute correlation of the embeddings with MOE descriptors but did not delve into the direction of correlation of these embeddings.

**Figure 4:**
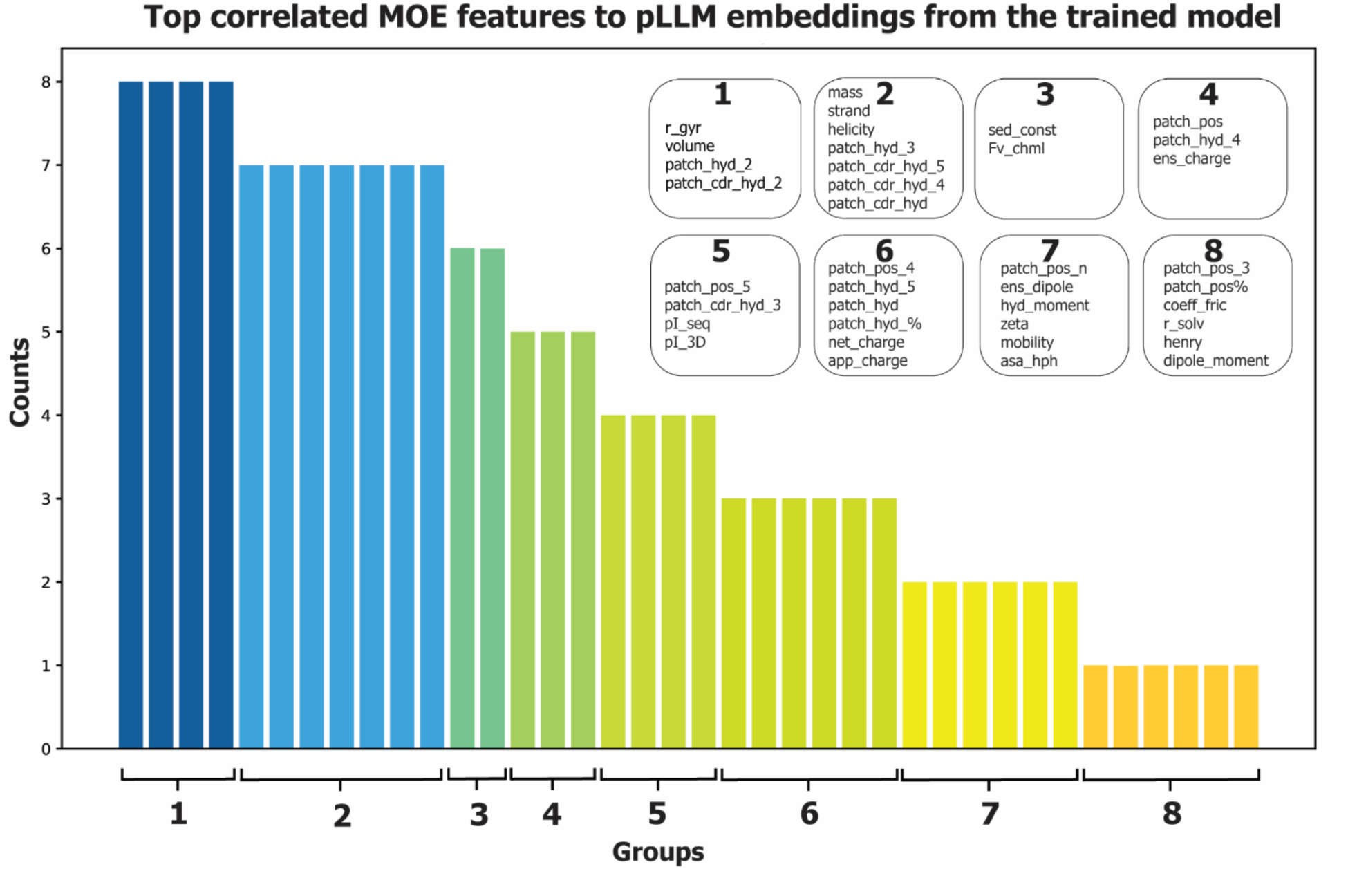
Top-ranked embeddings were interpreted based on their correlation with MOE features. Groups were defined by the frequency of MOE features appearing in the top 10 correlated features with pLLM embeddings. Group 1 includes features such as r_gyr, volume, patch_hyd_2, and patch_cdr_hyd_2, which appear 8 times, while Group 8 includes features like patch_pos_3, patch_pos%, coeff_fric, r_solv, henry, and dipole_moment, each occurring only once.

Hydrophobic patches on proteins, particularly within the complementarity-determining regions (CDRs) of antibodies, significantly influence protein viscosity by promoting self-association and aggregation. These solvent-exposed patches drive the formation of larger protein complexes, hindering molecular movement and increasing solution viscosity^14,17^. Similarly, charge asymmetry in the variable fragment (Fv) region of antibodies impacts protein viscosity at high concentrations by influencing intermolecular interactions and self-association tendencies. Studies have shown that viscosity correlates with both the net charge and charge asymmetry of the Fv region, with more positively charged variable domains generally exhibiting lower viscosity ^5,18^.

The impact of positive charge patches in the Fv region on protein viscosity is particularly significant at high concentrations. These patches tend to repel each other, reducing protein-protein interactions and aggregation, thus lowering solution viscosity. Conversely, negatively charged patches can attract positively charged regions on other antibodies, promoting aggregation and increasing viscosity^18^. Computational tools, such as the spatial charge map (SCM), have been developed to predict viscosity based on the distribution of electrostatic charges on the Fv surface, further highlighting the critical role of charge distribution in determining antibody solution viscosity^1^.

Recent research has also identified the dipole moment of monoclonal antibodies (mAbs) as a critical factor influencing the viscosity of concentrated antibody solutions. MAbs with larger dipole moments exhibit stronger dipole-dipole interactions, leading to increased protein-protein associations and higher viscosity in concentrated solutions^19^. This relationship becomes more pronounced at higher antibody concentrations, where the probability of dipole-dipole interactions between mAb molecules rises significantly. The spatial distribution of charges on the antibody surface, particularly in the Fab region, contributes to the overall dipole moment and plays a crucial role in determining viscosity behavior.

Our study has revealed several descriptors among the 38 identified features that, to our knowledge, have not been previously directly correlated with viscosity in existing literature. These factors, including helicity, mass, volume, and radius of gyration, are known to influence protein self-association and aggregation, but their explicit impact on viscosity has not been thoroughly explored. Our findings confirm the significant role these factors play in determining protein viscosity. The helicity of proteins, particularly the α-helical content, could influence viscosity through its effect on structural stability. Proteins with high α-helical content generally exhibit greater structural integrity, which can prevent partial unfolding - a precursor to aggregation^20^. By maintaining their structure, these proteins are less likely to expose hydrophobic regions that drive aggregation, resulting in lower viscosity at high concentrations.

The physical characteristics of monoclonal antibodies (mAbs), including their mass, volume, and radius of gyration (Rg), could also significantly impact solution viscosity, especially at high concentrations. The substantial molecular size of mAbs (∼150 kDa) leads to considerable excluded volume effects, reducing the available space for molecular movement and consequently increasing viscosity^21,22^. The Rg, which quantifies the spatial extent of the mAb, directly influences its hydrodynamic properties and solution behavior. Larger Rg values typically correlate with higher viscosities due to increased intermolecular interactions and reduced mobility. As mAb concentration increases, these effects become more pronounced, leading to an exponential rise in viscosity^23^. Our findings not only confirm the importance of previously recognized factors in determining protein viscosity but also highlight the significance of these additional parameters. This comprehensive understanding of the multifaceted influences on protein viscosity provides valuable insights for the development and optimization of therapeutic protein formulations, particularly in achieving desired viscosity profiles for high-concentration antibody solutions.

Direct correlation of MOE features with viscosity also yielded some useful insights. For descriptors such as Fv_chml, patch_hyd, pI, patch_pos charge, mass and volume, there is a high degree of correlation with the value of viscosity (|Spearman| >0.5) and statistical significance between the descriptor values for high and low viscous samples (using a cutoff of 20cP) (**SI Fig. 4**).

## 6 Conclusions, challenges and future directions

This study presents the first multi-modal machine learning model that integrates sequence and structural-level features from Molecular Operating Environment (MOE), and sequence-level embeddings from protein-specific language models. The information-rich embeddings from protein language models capture previously unrecognized, non-linear dependencies between sequence information and biophysical properties. When combined with molecular descriptors from MOE, this approach results in improved model performance for both regression and classification tasks. Despite these advancements, several limitations and opportunities for future research remain. Firstly, the viscosity of therapeutic proteins in formulation buffers is influenced by factors not fully accounted for in this model, such as buffer pH and the presence of excipients like salts and sugars. Secondly, the molecular descriptors used in this study are exclusively one-dimensional. Incorporating two-and three-dimensional surface descriptors, such as charge and hydrophobicity surface maps, could potentially capture additional dependencies and further enhance model performance. Lastly, while this study utilizes sequence-level embeddings, future work could explore the use of residue-level embeddings to increase the model’s sensitivity to viscosity changes resulting from single residue mutations.

In conclusion, this novel multi-modal approach represents a significant step forward in predicting therapeutic protein viscosity, but also highlights areas for continued refinement and expansion of the model to address complex protein-environment interactions.

## Supplementary Information

**Supplementary Fig.1.**
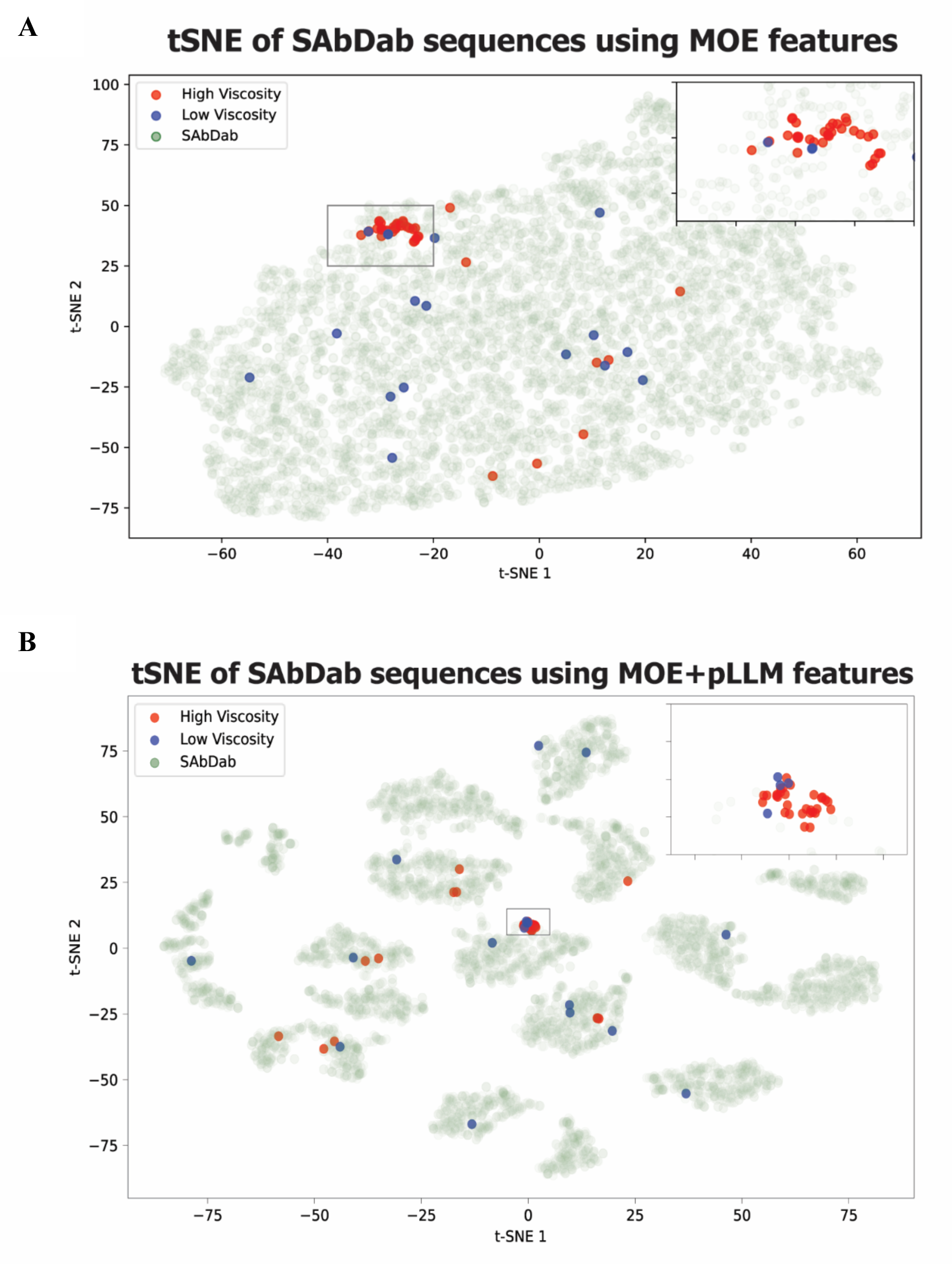
**(A,B):** t-SNE plot for all the SAbDab (6577 Ab) sequences overlaid with 59 (PDGF38+Ab21) sequences using A) 90 MOE features only and B)2909 (MOE+pLLM) features.

**Supplementary Table 1:** Isotype, HC_V gene, LC_V gene, cell line and experimental viscosity information for 15 in-house sequences.

| Sample ID | Isotype | LC | HC_V.Gene | LC_V.Gene | Cell Line | Measured Viscosity (cP)@150mg/ml |
| --- | --- | --- | --- | --- | --- | --- |
| GS-mAb1 | hIgG1 | kappa | IGHV3-30*01 | IGKV2-28*01 | CHO | 4.1 |
| GS-mAb2 | hIgG1 | Kappa | IGHV3-74*01 | IGKV1-33*01 | CHO | 2.8 |
| GS-mAb3 | hIgG1 | Kappa | IGHV3-48*03 | IGKV1-39*01 | Expi293 | 4.9 |
| GS-mAb4 | hIgG1 | Kappa | IGHV1-2*02 | IGKV2-30*02 | Expi293 | 7.9 |
| GS-mAb5 | hIgG1 | Kappa | IGHV2-5*08 | IGKV1-5*01 | CHO | 3.8 |
| GS-mAb6 | hIgG1 | Kappa | IGHV1-3*04 | IGKV1-NL1*01 | Expi293 | 4.1 |
| GS-mAb7 | hIgG1 | Kappa | IGHV7-4-1*02 | IGKV1-NL1*01 | CHO | 8 |
| GS-mAb8 | hIgG1 | Kappa | IGHV3-11*01 | IGKV4-1*01 | ExpiCHO | 4.7 |
| GS-mAb9 | hIgG1 | Kappa | IGHV1-46*01 | IGKV1-16*01 | Expi293 | 2 |
| GS-mAb10 | hIgG1 | Kappa | IGHV2-70*19 | IGKV1-5*01 | Expi293 | 3.7 |
| GS-mAb11 | hIgG1 | Kappa | IGHV1-2*06 | IGKV1-33*01 | Expi293 | 22.6 |
| GS-mAb12 | hIgG1 | lambda | IGHV1-2*06 | IGLV2-23*02 | Expi293 | 20 |
| GS-mAb13 | hIgG1 | Kappa | IGHV1-46*01 | IGKV1-NL1*01 | Expi293 | 9 |
| GS-mAb14 | hIgG1 | Kappa | IGHV3-23*04 | IGKV3/OR2-268*01 | Expi293 | 6.5 |
| GS-mAb15 | hIgG1 | Kappa | IGHV7-4-1*02 | IGKV1-33*01 | Expi293 | 5 |

**Supplementary Figure 2:**
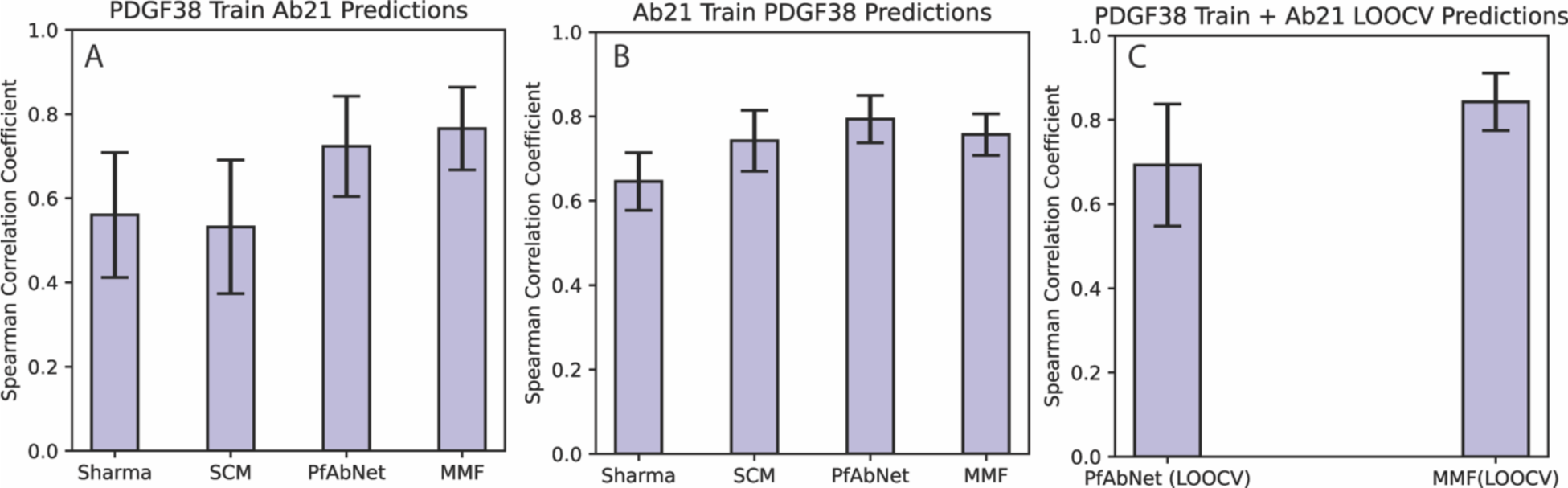
Bootstrapping analysis (n=500) followed by t-test to evalu<u>in</u>ate the significance in spearman rank order correlation between various models. A) PDGF38 as the training set and Ab21 as the test set, B) Ab21 as the train set and PDGF38 as the test set and C) Leave one out cross validation (LOOCV) for PfAbNet vs MMF.

**Supplementary Table 2:** Pairwise t-test comparison of the bootstrap results.

| Model Comparison | Pairwise t-test p value (Ab21 Predictions) | Pairwise t-test p value (PDGF38 Predictions) |
| --- | --- | --- |
| Sharma vs SCM | 3.81e-03 | 4.05e-83 |
| Sharma vs PfAbNet | 2.32e-66 | 3.20e-182 |
| Sharma vs MMF | 4.86e-101 | 3.46e-115 |
| SCM vs PfAbNet | 9.89e-81 | 4.47e-35 |
| SCM vs MMF | 7.15e-114 | 9.70e-03 |
| PfAbNet vs MMF | 9.61e-07 | 1.16e-31 |

**Supplementary Table 3A:**
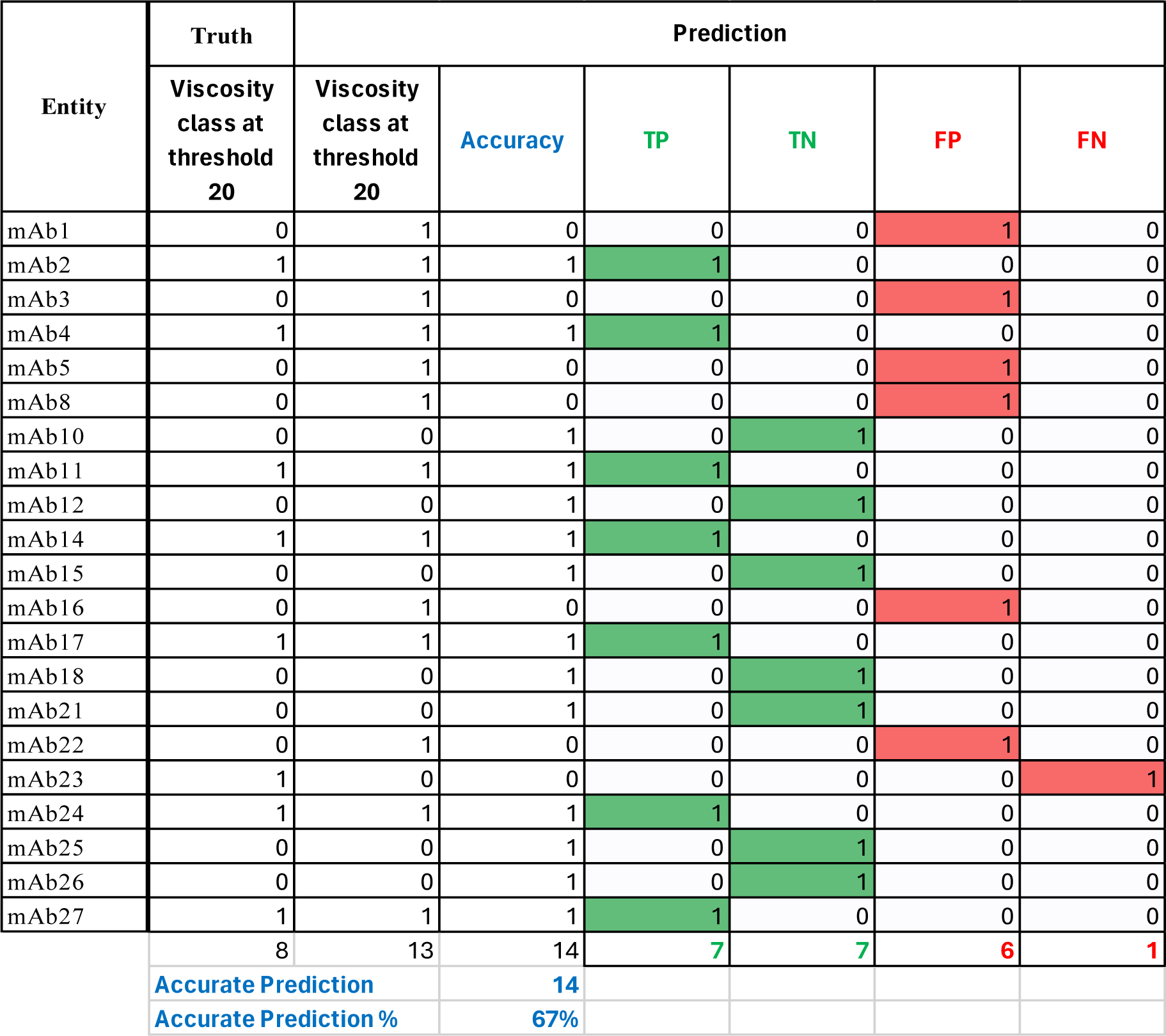
Classification accuracy of PfAbNet at practical viscosity cutoff value of 20cP.

**Supplementary Table 3B:** Classification accuracy of MMF at practical viscosity cutoff value of 20cP.

| Entity | Truth | Prediction |  |  |  |  |  |
| --- | --- | --- | --- | --- | --- | --- | --- |
|  | Viscosity class at threshold 20 | Viscosity class at threshold 20 | Accuracy | TP | TN | FP | FN |
| mAb1 | 0 | 0 | 1 | 0 | 1 | 0 | 0 |
| mAb2 | 1 | 1 | 1 | 1 | 0 | 0 | 0 |
| mAb3 | 0 | 0 | 1 | 0 | 1 | 0 | 0 |
| mAb4 | 1 | 1 | 1 | 1 | 0 | 0 | 0 |
| mAb5 | 0 | 0 | 1 | 0 | 1 | 0 | 0 |
| mAb8 | 0 | 0 | 1 | 0 | 1 | 0 | 0 |
| mAb10 | 0 | 0 | 1 | 0 | 1 | 0 | 0 |
| mAb11 | 1 | 1 | 1 | 1 | 0 | 0 | 0 |
| mAb12 | 0 | 0 | 1 | 0 | 1 | 0 | 0 |
| mAb14 | 1 | 0 | 0 | 0 | 0 | 0 | 1 |
| mAb15 | 0 | 1 | 0 | 0 | 0 | 1 | 0 |
| mAb16 | 0 | 0 | 1 | 0 | 1 | 0 | 0 |
| mAb17 | 1 | 1 | 1 | 1 | 0 | 0 | 0 |
| mAb18 | 0 | 0 | 1 | 0 | 1 | 0 | 0 |
| mAb21 | 0 | 0 | 1 | 0 | 1 | 0 | 0 |
| mAb22 | 0 | 0 | 1 | 0 | 1 | 0 | 0 |
| mAb23 | 1 | 1 | 1 | 1 | 0 | 0 | 0 |
| mAb24 | 1 | 1 | 1 | 1 | 0 | 0 | 0 |
| mAb25 | 0 | 0 | 1 | 0 | 1 | 0 | 0 |
| mAb26 | 0 | 0 | 1 | 0 | 1 | 0 | 0 |
| mAb27 | 1 | 1 | 1 | 1 | 0 | 0 | 0 |
|  |  |  |  | 7 | 12 | 1 | 1 |
| Accurate Prediction |  |  | 19 |  |  |  |  |
| Accurate Prediction % |  |  | 90% |  |  |  |  |

**Supplementary Figure 3:**
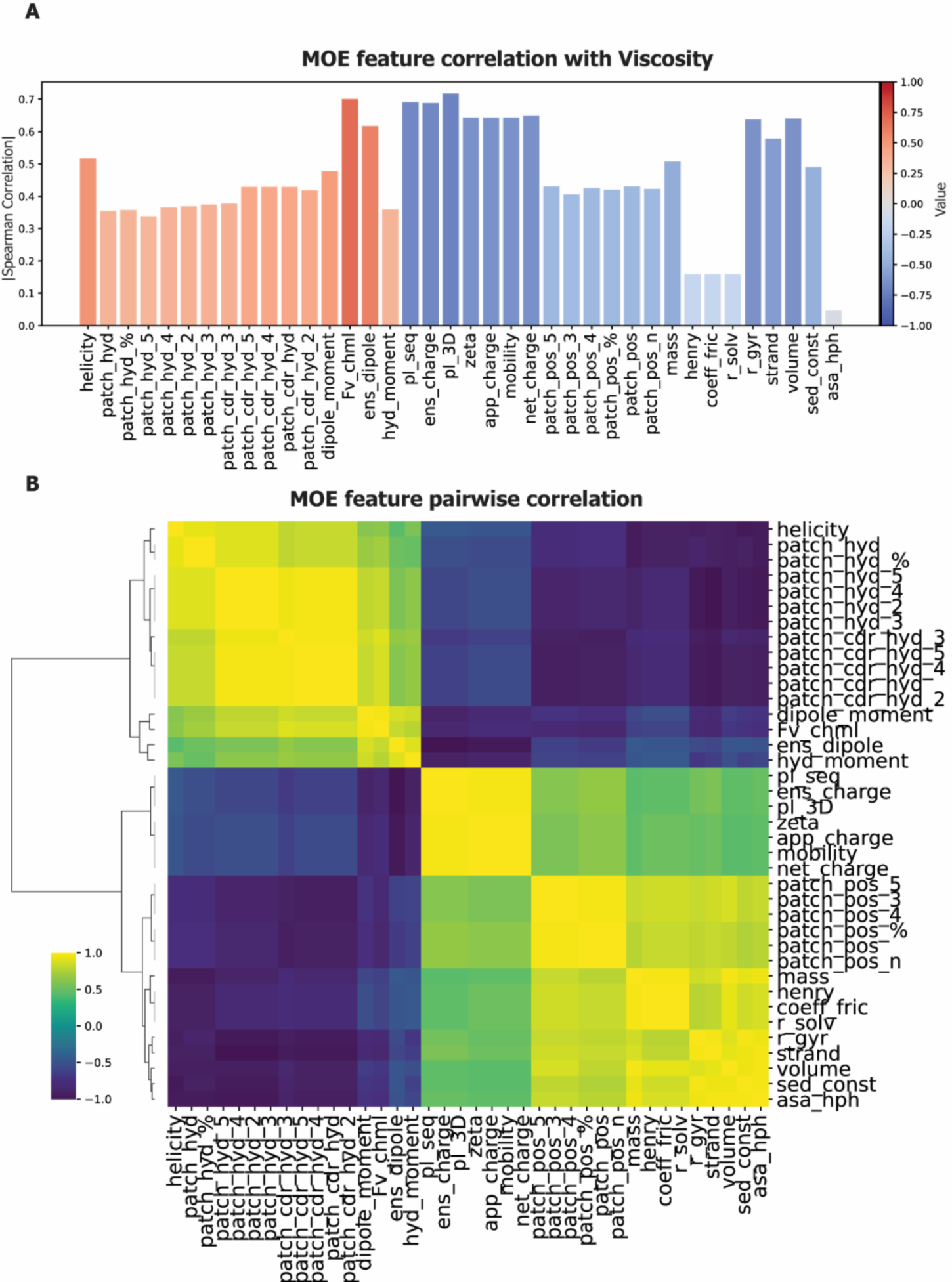
A) MOE feature correlation with measured viscosity value for the 59 mAbs (PDGF38+Ab21). B) Pairwise correlation among MOE features.

**Supplementary Figure 4:**
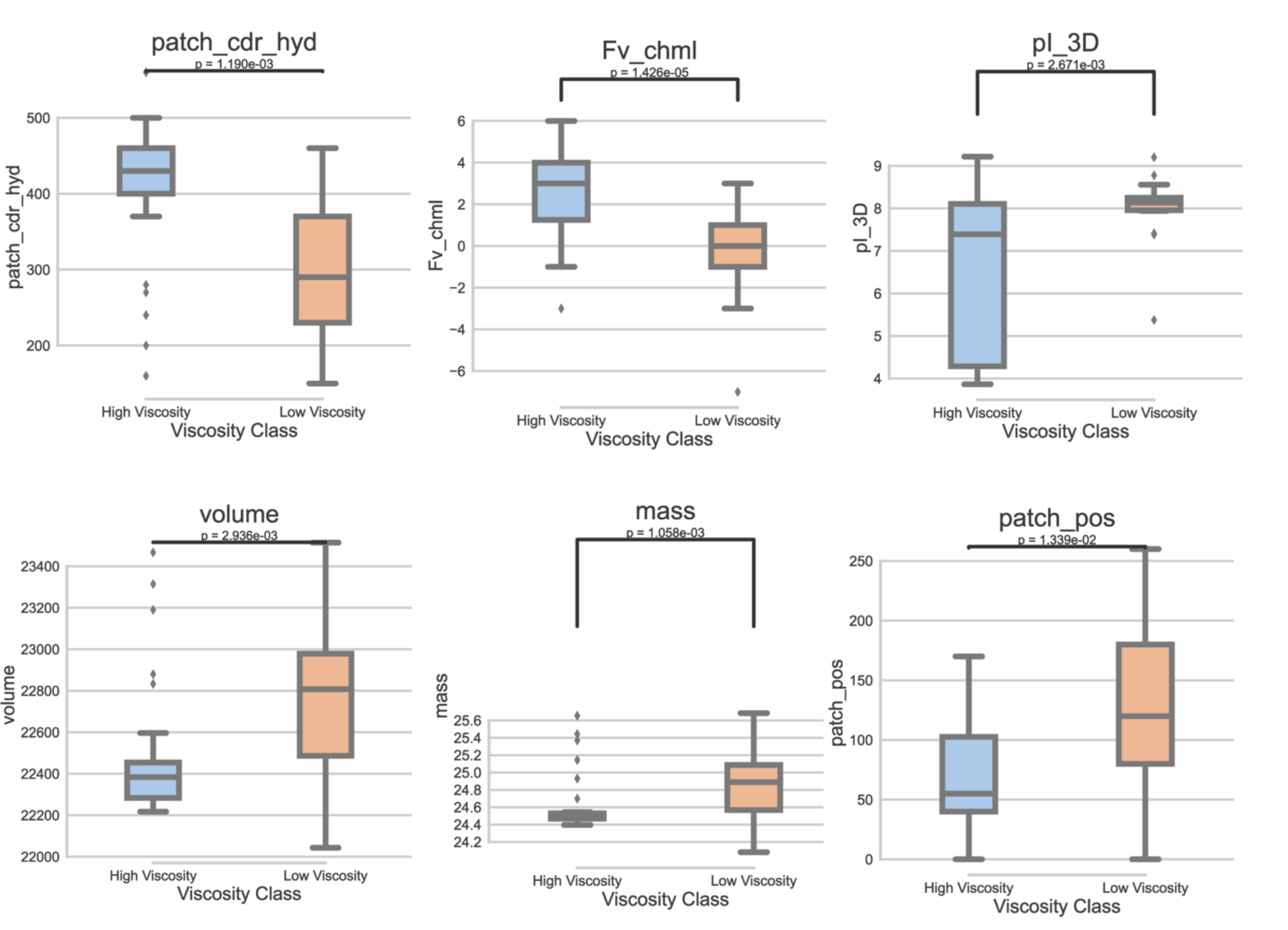
Box plots comparing the distribution of 6 different MOE properties for low and high viscous samples from the 59 mAb sequences used in the training and testing of MMF.

## Appendix I *MOE_Property_Description.xlsx*: List of all the 38 MOE properties correlated with pLLM embeddings and their description.

## Notes

### Competing Interest Statement

The authors have declared no competing interest.

